# Freeze tolerance of a beneficial lady beetle, *Hippodamia convergens*

**DOI:** 10.64898/2026.08.13.744689

**Authors:** Haley E. Ehler, Teresa A. Keenan, Lousie E. Evans, Madelyn M.R. Weisshaar, Emma L. Barrett, Liam J. Kirkham, Jalen K. Macfarlane, Anna B. Silver, María F. Alas Romero, Brenna R. Alford, Ariana M. Ayala Rosero, Ryan P. Clancy, Sophie D. Fraser, Gracyn M. Glennie, Kiersten E. D. Hooper, Kiersten E. MacGrath, Jaylene M. Nauss, Lauren A. Pictou, Maria K. Putnam, Zoe M. Sturmy, Jennifer C. Perry, Jantina Toxopeus

## Abstract

The convergent lady beetle *Hippodamia convergens* is widespread in the Americas and considered an important beneficial insect due to its use in pest control. Early work on this species characterized the beetles as freeze-avoidant (freeze-intolerant), suggesting they survive low winter temperatures by physiologically preventing ice formation to temperatures as low as −15°C. Here, we show that *H. convergens* can be freeze-tolerant if ice formation occurs at relatively high temperatures. There was 100% survival following inoculative freezing at −0.5°C and exposure to −3°C for 20 hours, as well as freezing that spontaneously occurred in fed beetles exposed to −4°C for 4 hours. Males exposed to 0°C or −4°C for 4 hours had similar mating behaviours (latency to first mating, copulation duration) as control beetles exposed to 4°C, although sample sizes were too small to determine whether freezing itself had an effect on these behaviours. Several putative cryoprotectants were detected in fat body tissue of *H. convergens*: glycerol, proline, trehalose, and *myo*-inositol – in order of abundance. Exposure to −3°C for 20 hours, whether frozen or unfrozen, did not statistically affect cryoprotectant accumulation, although there was a trend towards increased glycerol following freezing. This study is the first to describe inoculative freeze tolerance in a lady beetle and establishes a baseline for future studies that examine the mechanisms underlying this freeze tolerance and the effects of freezing on reproductive behaviour.

**Graphical abstract:** 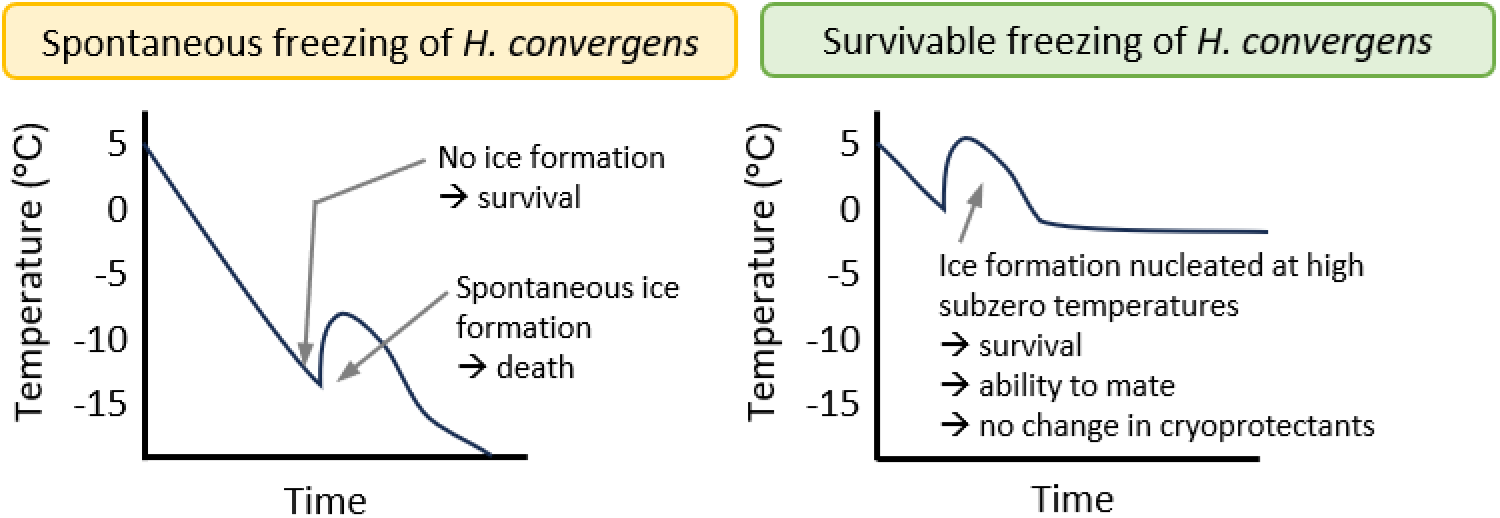

- *Hippodamia convergens* can tolerate freezing at high subzero temperatures
- Mating behaviour of males is normal after chilling or freezing
- Cryoprotectant accumulation may support overwintering survival

## Introduction

Low temperatures can cause strong selective pressures on organisms, either directly by causing mortality, or indirectly via sublethal effects on fitness. For example, subzero temperatures can cause freezing of internal body fluids, which is lethal for most insect species (Toxopeus and Sinclair, 2018). Mild cold (e.g., temperatures near 0°C) can also impair organismal function and lead to lethal chilling injuries (Overgaard and MacMillan, 2017). Insects that survive low temperatures may have reduced fitness due to negative effects of chilling or freezing on reproduction (Koštál et al., 2019; Marshall and Sinclair, 2018, 2011). However, exposure to mild low temperatures can also improve insect survival and fitness by increasing tolerance to subsequent stressful conditions (Teets and Denlinger, 2013). It is important to understand how insects are affected by, and respond to, low temperatures to be able to predict their overwintering survival, particularly as winters become warmer and more variable due to climate change (Marshall et al., 2020; Williams et al., 2015).

Insects that survive low temperatures are usually classified with a cold tolerance strategy of freeze tolerance or freeze avoidance (Sinclair et al., 2015), and each strategy is associated with different cold tolerance mechanisms. As an insect is cooled below 0°C, ice formation can begin at a temperature known as the supercooling point (Sinclair et al., 2015), which varies within and among species. Freeze-tolerant insects survive this spontaneous freezing at the supercooling point, while freeze-avoidant insects do not (Toxopeus and Sinclair, 2018). However, Rozsypal and Koštál (2018) have shown that freeze-avoidant insects may survive internal ice formation that is initiated at a relatively high subzero temperature (e.g., −2.5°C) via contact with external ice, a process known as inoculative freezing. In addition, some insects are characterized as partially freeze-tolerant, and can tolerate some internal ice formation at their supercooling point for short periods of time, as seen in the harlequin lady beetle *Harmonia axyridis* (Koch et al., 2004; Sinclair, 1999). Freeze-avoidant insects such as the eastern spruce budworm *Choristoneura fumiferana* typically survive low subzero temperatures by decreasing their supercooling point (to below −30°C!) using a combination of cryoprotectant molecules (e.g., glycerol) and antifreeze proteins (Han and Bauce, 1995; Lee, 2010; Tyshenko et al., 1997). Freeze-tolerant insects such as the spring field cricket *Gryllus veletis* are hypothesized to use a variety of mechanisms to survive freezing, including accumulating cryoprotectants to protect cellular components and ice-nucleating agents to control ice formation (Toxopeus et al., 2019a, 2019b; Toxopeus and Sinclair, 2018). In temperate regions, many insects increase their cold tolerance seasonally, for example accumulating cryoprotectants prior to the onset of winter (Sinclair et al., 2015; Toxopeus et al., 2019a).

While surviving low temperatures is important for insects in temperate climates, being able to reproduce post-winter is key for fitness. Winter conditions are expected to negatively affect reproduction (Meuti et al., 2024), and even short cold exposures (e.g., a few hours at 0°C) cause decreased reproductive success in warm-adapted species such as the house cricket *Acheta domesticus* (Chipchase et al., 2021) and the red cotton bug *Dysdercus koenigii* (Sarmad et al., 2020). Exposure to winter conditions (November to March) in the field causes a delay and decrease in the number of eggs laid by female willow leaf beetles, *Plagiodera versicolora* (Yan et al., 2025). However, natural overwintering better supports reproduction (higher fecundity, fertility) than prolonged cold storage (e.g., 5°C for 18 weeks) in green lacewings, *Chrysoperla carnea* (Chang et al., 1996). Cold exposure can also affect mating behaviour, with *Drosophila melanogaster* males showing longer latency to first mating after a short cold shock (1 hour at −5°C; Singh et al., 2016), a trait that can indicate lower male fitness (Bista and Omkar, 2015). Latency to mating can shorten over time post-cold shock (Singh et al., 2016), indicating likely recovery from chilling injuries.

The convergent lady beetle *Hippodamia convergens* Guérin-Méneville (Coleoptera: Coccinellidae) is widespread in North and South America (Obrycki et al., 2001). As predators of aphids, these lady beetles are a popular choice for pest control (e.g., Flint and Dreistadt, 2005; Rashed et al., 2020). In the temperate regions of their range, *H. convergens* overwinter as adults in reproductive diapause (Michaud and Qureshi, 2006, 2005; Nadeau et al., 2022; Obrycki et al., 2018), suppressing development of gonads to ensure sufficient energy reserves can be directed toward survival of adverse conditions (Hahn and Denlinger, 2011). In addition, *H. convergens* often overwinter in aggregations that may help them conserve energy (Simpson and Welborn, 1975; Szejner-Sigal and Williams, 2022). Prior work has shown that cold-acclimated *H. convergens* freeze spontaneously at approximately −15°C and do not survive this freezing (Bennett and Lee, 1989; Lee, 1980). However, freezing can be simulated at higher temperatures (e.g., −3 to −4°C) by ingestion of ice-nucleating *Pseudomonas syringae* or *Erwinia herbicola* bacteria (Strong-Gunderson et al., 1990), or application of *P. syringae* or *Fusarium avenaceum* (fungus) to beetle thoracic spiracles (Steigerwald et al., 1995). However, survival was not described following these freeze treatments (Steigerwald et al., 1995; Strong-Gunderson et al., 1990). To our knowledge, there has not been any work on the effect of cold exposure on reproduction in *H. convergens*.

In this study, we characterized the cold and freeze tolerance of *H. convergens* with three main goals. First, we determined whether beetles in diapause could survive freezing under a variety of conditions, including inoculative freezing at mild subzero temperatures. We predicted that these insects would be freeze-avoidant (Bennett and Lee, 1989; Lee, 1980) but had the potential to survive freezing induced at higher temperatures (Rozsypal and Koštál, 2018). Second, we examined whether post-diapause male beetles could still mate following exposure to mild low temperatures. If low temperatures caused chilling or freezing injury, we predicted this would decrease the proportion of males that successfully copulated and increase the latency to first mating (Bista and Omkar, 2015; Singh et al., 2016). Third, we measured concentrations of some common insect cryoprotectants in diapause beetles. We predicted these cryoprotectant concentrations would increase following exposure to chilling or freezing at mild low temperatures (Li et al., 2020; Toxopeus et al., 2016) to facilitate survival of future low temperature exposures.

## Materials and Methods

### Insect rearing and acclimation

*Hippodamia convergens* (Costco, Dartmouth NS, Canada or NaturesArt, Toronto ON, Canada) were shipped to St. Francis Xavier University (Antigonish NS, Canada) under ambient conditions in September - October 2024 and 2025 (diapause beetles) or January 2024 (post-diapause beetles). Upon arrival, they were separated by sex and transferred in small groups (20-30 beetles) to Petri dishes with food (frozen Nutrimac - *Ephestia kuehniella* eggs; Plant Products, Leamington ON, Canada) and water-soaked cotton balls. Beetles were kept at room temperature under natural light conditions for up to 3 weeks before being transferred to 4°C and darkness in a fridge for up to 4 weeks prior to use in the experiments described below.

Beetles that arrived in September or October were confirmed to be non-feeding, and likely in diapause. A subset of these beetles were dissected to confirm large accumulation of fat stores (abdomen primarily filled with fat body tissue) and reduced gonads, consistent with a diapause phenotype (Michaud and Qureshi, 2005). Beetles that arrived in January did engage in feeding when at room temperature, indicating they were likely in a post-diapause state (Michaud and Qureshi, 2006).

### Assessing cold tolerance

To determine the supercooling point (temperature of spontaneous freezing) of diapause beetles, 6 males and 6 females were each placed in a 1.7 ml tube in contact with a thermocouple and cooled by a programmable recirculating chiller (Adams et al., 2025; Lemay et al., 2024; Lopez Pedersen et al., 2026). The Arctic A25 chiller (Thermo Fisher Scientific, Toronto, ON, Canada) contained 50% v/v propylene glycol that was circulated to a custom insulated aluminum block containing the tubes. The tubes were cooled from 4°C to −20°C at a rate of −0.25°C min^-1^. Temperature was recorded via T-type copper-constantan thermocouples (Omega Engineering, Norwalk, CT, USA) interfaced with Picolog v. 6.2.8 software via a TC-8 unit (Pico Technology, Cambridgeshire, UK). The supercooling point was the lowest temperature recorded prior to an exotherm - an increase in temperature associated with internal ice formation (Sinclair et al., 2015).

To determine the cold tolerance strategy (as defined by Sinclair et al., 2015) of diapause beetles, groups of 21 male or female beetles were cooled to a temperature (c. −15°C) at which approximately half of the beetles froze and half remained unfrozen, followed by assessment of survival (Adams et al., 2025; Lemay et al., 2024; Lopez Pedersen et al., 2026). Beetles were placed in tubes and cooled as described for the supercooling point measurement, followed by immediate transfer to 4°C once half the beetles had frozen. Beetles were defined as frozen if they reached their supercooling point and experienced ice formation for at least 5 minutes. Beetles were defined as unfrozen (supercooled) if they did not reach their supercooling point. A small portion of beetles were defined as partially frozen – they reached their supercooling point, but ice formation only occurred for 1 – 5 seconds before beetles were transfer to 4°C. In addition to beetles that were cooled in the recirculating chiller, 8 females and males were included as handling controls that were placed in tubes but remained at 4°C while the experimental beetles were being cooled. Survival (ability to move) of all beetles was assessed 48 hours after the cooled beetles were transferred to 4°C. Beetles were defined as freeze-tolerant if they survived freezing, freeze-avoidant if they survived supercooling but not freezing, or chill-susceptible (not cold-tolerant) if survival was low for all cooled beetles (Sinclair et al., 2015).

To determine whether diapause beetles could survive inoculative freezing at temperatures above their supercooling point, females and males were exposed to −3, −5, or −8°C for approximately 1 day, followed by return to 4°C for 48 hours and survival assessment. We followed a modified version of the protocol used by Rozsypal and Koštál (2018) to cause inoculative freezing. Prior to the cold exposure, beetles were each surrounded by a small square (4 - 5 cm^2^) of wet paper towel and placed in a 1.7 mL tube in contact with a thermocouple. These tubes were transferred to the aluminum block pre-chilled to −0.5°C. A small piece of ice was added to half of the tubes to stimulate freezing of the water in the paper towel, placing the beetles in direct contact with ice. After 20-30 minutes at −0.5°C, the beetles were cooled to their target temperature at −0.25°C min^-1^, held at their target temperature for 20 hours, and then warmed to 4°C at 0.25°C min^-1^. Some paper towel squares froze (detectable as a large exotherm) even when ice was not added, and the proportion of beetles in contact with frozen paper towel that froze (a smaller exotherm) increased at lower temperatures (Table 1). Similar to the cold tolerance strategy experiment above, we included handling controls at 4°C in parallel with each low temperature exposure. We did not conduct a dedicated inoculative freezing experiment with post-diapause beetles. However, some post-diapause beetles spontaneously froze when exposed to −4°C prior to the mating behaviour assays described below, which we expand on in the Results.

**Table 1.** Effectiveness of inoculative freezing manipulation in diapause beetles. Beetles encased in wet paper towel were cooled to a target temperature (−3, −5, or −8°C) for 20 hours. A small piece of ice was added to half of the tubes at −0.5°C to cause the paper towel, and subsequently the beetle, to freeze. The proportions of wet paper towel and beetles that froze are shown.

| Temperature | Ice added |  | Ice not added |  | Beetles in contact with ice that froze<sup>c</sup> |
| --- | --- | --- | --- | --- | --- |
|  | Paper towel froze | Beetle froze<sup>a</sup> | Paper towel froze<sup>b</sup> | Beetle froze<sup>a</sup> |  |
| -3°C | 30/32 | 12/32 | 5/32 | 3/32 | 15/35 (43%) |
| -5°C | 8/8 | 3/8 | 6/8 | 5/8 | 8/14 (57%) |
| -8°C | 8/8 | 7/8 | 7/8 | 6/8 | 13/15 (87%) |

<sup>a</sup> All beetles that froze were in contact with paper towel that froze (i.e., no spontaneous freezing).
<sup>b</sup> Freezing of paper towels could be caused by exposure to the target temperature rather than the presence of a piece of ice in the tube.
<sup>c</sup> Number of beetles that froze / Number of paper towel squares that froze

### Mating behaviour assays

To determine if cold exposure affected mating behaviour in post-diapause beetles, males were exposed to one of three treatments using the fridge and recirculating chiller. Following 3 weeks at room temperature with food and water, control males (*N* = 8) were kept in the fridge at 4°C for one week. The other two groups were kept at 4°C for 1 week and then transferred in 1.7 ml tubes into the pre-cooled aluminum block attached to the recirculating chiller at 0°C (*N* = 15) or −4°C (*N* = 7) for 4 hours. The treatment temperature was confirmed using thermocouples in each tube. Unexpectedly, 5 of the 7 males exposed to −4°C froze, as determined by the presence of an exotherm, but all survived. After the temperature treatment, beetles were returned to 4°C in individual petri dishes with food and water for 1 day prior to mating behaviour assays.

For the mating behaviour assays, each cold-exposed male was paired with a female that had been kept under control conditions and then observed on three consecutive days. On the first mating day, each female was added to a male’s small petri dish after 1 hour at room temperature, and mating was observed for 8 hours before each pair was separated and returned to 4°C. On the second and third mating days, a similar protocol was followed but the observation period was 4 hours. During observation, each pair was observed every 5 minutes. Latency (time to first mating) and copulation duration were recorded on each day. Mating was defined as when the male copulatory organ was inside the female. Any copulations less than 30 minutes long were defined as failed mating attempts due to insufficient time for sperm transfer. Males and females were moved into separate petri dishes once a mating event (successful or failed) ended.

### Spectrophotometric assays

To determine whether diapause beetles accumulate cryoprotectants, and whether this accumulation is affected by low temperature exposure, we conducted several spectrophotometric assays on fat body tissue. Fat body was dissected from beetles that had been exposed to −3°C for 20 hours and frozen (*N* = 7 females, 7 males) or remained unfrozen (*N* = 9 females, 8 males), along with handling controls that remained at 4°C (*N* = 8 females, 8 males), followed by recovery as described for the inoculative freezing experiments. This fat body was dissected 48 hours post-cold exposure, flash-frozen in liquid nitrogen, and stored at −80°C until processed for spectrophotometric assays (Lopez Pedersen et al., 2026; Toxopeus et al., 2019a, 2016).

To prepare samples for spectrophotometric assays, each fat body sample was homogenized on ice in 100 µL Tris-buffered saline (TB; 5 mM Tris; 137 mM NaCl; 2.7 mM KCl, pH 6.6) using a plastic micropestle (Thermo Fisher). 8 µL of homogenate was aliquoted for assaying proline concentration. The remaining homogenate was heat-treated at 70°C for 10 minutes to denature any endogenous enzymes (Tennessen et al., 2014), followed by 3 minutes centrifugation at 20,000 × *g*. The supernatant was aliquoted into 1.7 ml tubes for a glycerol assay (8 µL), *myo*-inositol assay (50 µL), and trehalose assay (20 µL). In pilot studies, heat-treated homogenate was also used to measure mannitol (D-Mannitol Assay Kit; Megazyme, Bray, Ireland) and sorbitol (D-Sorbitol/Xylitol Assay Kit; Megazyme), but neither of these putative cryoprotectants were detectable.

Glycerol concentration was determined using the Glycerol GK Assay Kit (Megazyme), in which the homogenate was diluted 1/5 in TB, and absorbance at 340 nm was measured in each sample before and 7 minutes after the addition of a glycerokinase enzyme that (in combination with ADP-glucokinase and diaphorase) catalyzes the formation of NADH. End point absorbance was measured in triplicate in a 96-well plate using a SpectraMax iD3 spectrophotometer (Molecular Devices, San Jose, CA, USA). Glycerol concentration in each sample was determined from the delta absorbance value (absorbance after – before enzyme addition) compared to a standard curve from 0.013 to 0.20 mg/ml glycerol standards.

*myo*-Inositol concentration was determined using the *myo*-Inositol Assay Kit (Megazyme), in which *myo*-inositol dehydrogenase, diaphorase, and INT (iodonitrotetrazolium) are used to generate an orange product (INT-formazan) that absorbs light at 492 nm. End point absorbance was measured at 492 nm in triplicate in a 96-well plate using the iD3 spectrophotometer. *myo*-Inositol concentration in each sample was determined by comparison to a standard curve of 0.016 to 0.25 mg/ml *myo*-inositol.

Proline concentration was determined using the assay described by Carillo and Gibon (2011), in which proline was extracted from the homogenate (diluted 1/10) overnight in 40 % (v/v) ethanol, followed by a heat and acid-catalyzed reaction with ninhydrin solution (1% [w/v] ninhydrin, 60% [v/v] acetic acid, 20% [v/v] ethanol) at 95°C for 20 minutes. End point absorbance was measured in triplicate at 520 nm in a 96-well plate using the iD3 spectrophotometer. Proline concentration in each sample was determined by comparison to a standard curve of 0.03 to 0.5 mM proline.

Trehalose concentration was determined using the assay described by Tennessen et al. (2014) and modified by Toxopeus et al. (2019a), in which trehalose in the homogenate is digested by porcine trehalase (Sigma Aldrich, Toronto, ON, Canada) and then glucose concentration is determined using the Glucose Assay Reagent (SigmaAldrich), which uses hexokinase and G6P dehydrogenase to catalyze the formation of NADH. Each homogenate was divided into two tubes – one that included trehalase (glucose generated by digestion of trehalose) and one that did not (only “background” glucose present) – and diluted 1/10 prior to the glucose assay. End point absorbance of each digested and undigested sample was measured in triplicate at 340 nm in a 96-well plate using an iD3 spectrophotometer. Trehalose concentration in each sample was determined by comparing the delta absorbance (digested - undigested) to that of standard curves of digested and undigested 0.01 to 0.016 mg/ml trehalose.

### Statistical analyses

All statistical analyses were conducted in R v. 4.5.2 (R Core Team, 2026). To determine whether supercooling point differed between males and females, we used an ANCOVA with mass as a covariate. We used a G-test (McIntyre et al., 2023) to determine whether the proportion of surviving frozen and chilled beetles differed across temperatures in the inoculative freezing experiment. To compare reproductive behaviours among observations days, we focused only on the first 4 hours (out of 8 hours) of observation on day 1 to use a comparable time frame to the 4-hour observation periods on days 2 and 3. We used a G-test to determine whether the proportion of males that mated during the 4 hour observation periods post-cold exposure varied with temperature treatment and date of mating observation (day 1, 2, or 3). To determine the effect of cold exposure and observation date on latency and copulation duration during the 4-hour observation periods, we used two-way ANOVAs. We also compared latency and copulation duration across cold treatments during the full 8-hour observation period on mating day 1 using ANOVAs. To determine the effect of chilling or freezing at −3°C on the concentrations of putative cryoprotectants, we used ANOVAs - one each for glycerol, *myo*-inositol, proline, and trehalose.

## Results

### Diapause lady beetles do not tolerate spontaneous freezing at low subzero temperatures

Our initial characterization of lady beetle cold tolerance suggested that diapause beetles could supercool to avoid freezing, facilitating survival of low temperatures. Females and males spontaneously froze at mean temperatures of −16°C and −15.5°C, respectively (Figure 1). Neither mass nor sex significantly affected supercooling point (ANCOVA: Mass *F*_1,10_ = 0.117, *P* = 0.740; Sex *F*_1,10_ = 0.725, *P* = 0.417). The beetles did not survive spontaneous freezing but did survive brief exposures to −15°C if they remained unfrozen (Table 2), consistent with a freeze avoidance strategy. However, a small number of beetles partially froze in the cold tolerance strategy experiment and survived (Table 2), suggesting *H. convergens* may tolerate ice formation under specific circumstances.

**Figure 1.**
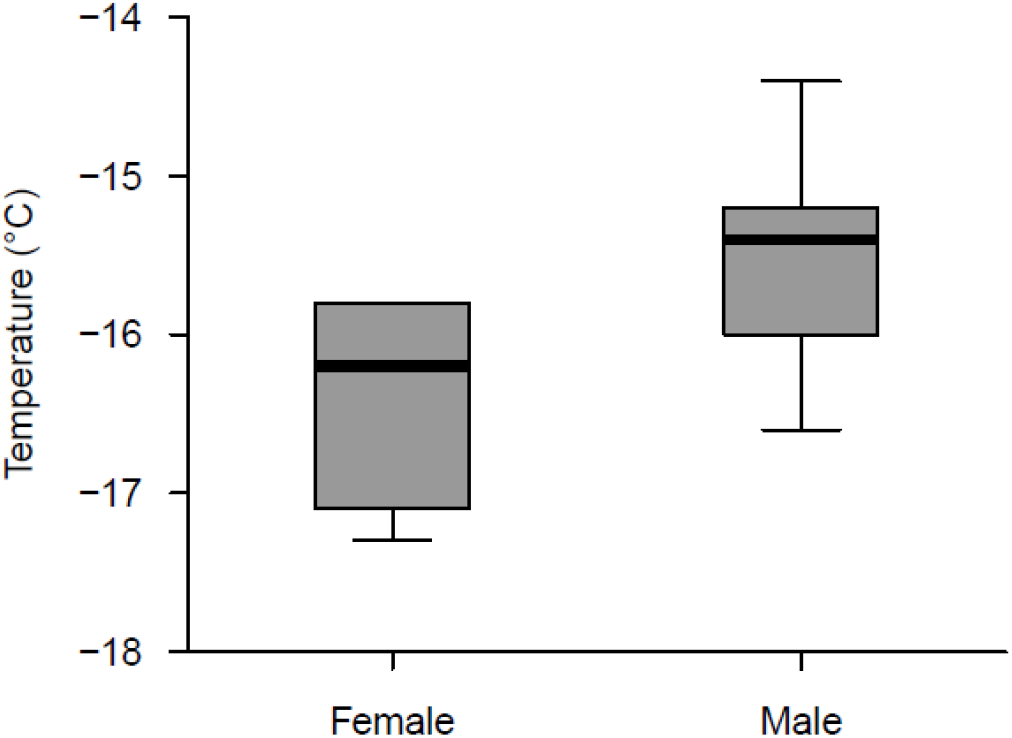
Temperature of spontaneous internal freezing (supercooling point) of male and female *Hippodamia convergens* in diapause. The thick central line is the median, with the lower and upper bounds of the box representing the first and third quartile, respectively. Whiskers extend to 1.5 times the interquartile range or the most extreme data point above/below the box, whichever is shortest.

**Table 2.** Survival of diapause *Hippodamia convergens* from the cold tolerance strategy experiment. Groups of beetles were cooled to a temperature (c. −15°C) at which half of them froze and half remained unfrozen, followed by 2 days recovery at 4°C.

|  | N survived/<br>N supercooled<sup>a</sup> | N survived/<br>N frozen<sup>b</sup> | N survived/<br>N partially frozen<sup>c</sup> | N survived/<br>N controls<sup>d</sup> |
| --- | --- | --- | --- | --- |
| Females | 11/12 | 0/8 | 1/1 | 8/10 |
| Males | 8/9 | 0/9 | 3/3 | 10/10 |

<sup>a</sup>Supercooled individuals never froze.
<sup>b</sup>Frozen individuals were kept in the chiller for at least 5 minutes after starting to freeze.
<sup>c</sup>Partially frozen individuals started to freeze 1 – 5 seconds prior to removal from the chiller.
<sup>d</sup>Controls were handled similarly to experimental beetles, but remained at 4°C.

### Diapause and post-diapause lady beetles tolerate freezing at high subzero temperatures

Contrary to our initial assessment of the beetles as freeze-avoidant, they were able to survive freezing when frozen at relatively high temperatures, even after diapause was complete. When diapause beetles were exposed to −3°C for 20 hours in contact with external ice, almost all beetles that froze were able to recover motility within 48 hours post-thaw, along with beetles that did not freeze (Figure 2A). However, beetles that were inoculatively frozen at lower temperatures (−5°C and −8°C) did not survive, while some that remained unfrozen (chilled) at these temperatures did survive (Figure 2A). Almost all (47/49) handling controls (kept at 4°C) survived for the duration of the experiment, so the death observed at −5°C and −8°C was driven by exposure to low temperatures and freezing rather than general beetle condition. All post-diapause beetles that froze or remained chilled during a 4-hour exposure to −4°C also survived (Figure 2B).

**Figure 2.**
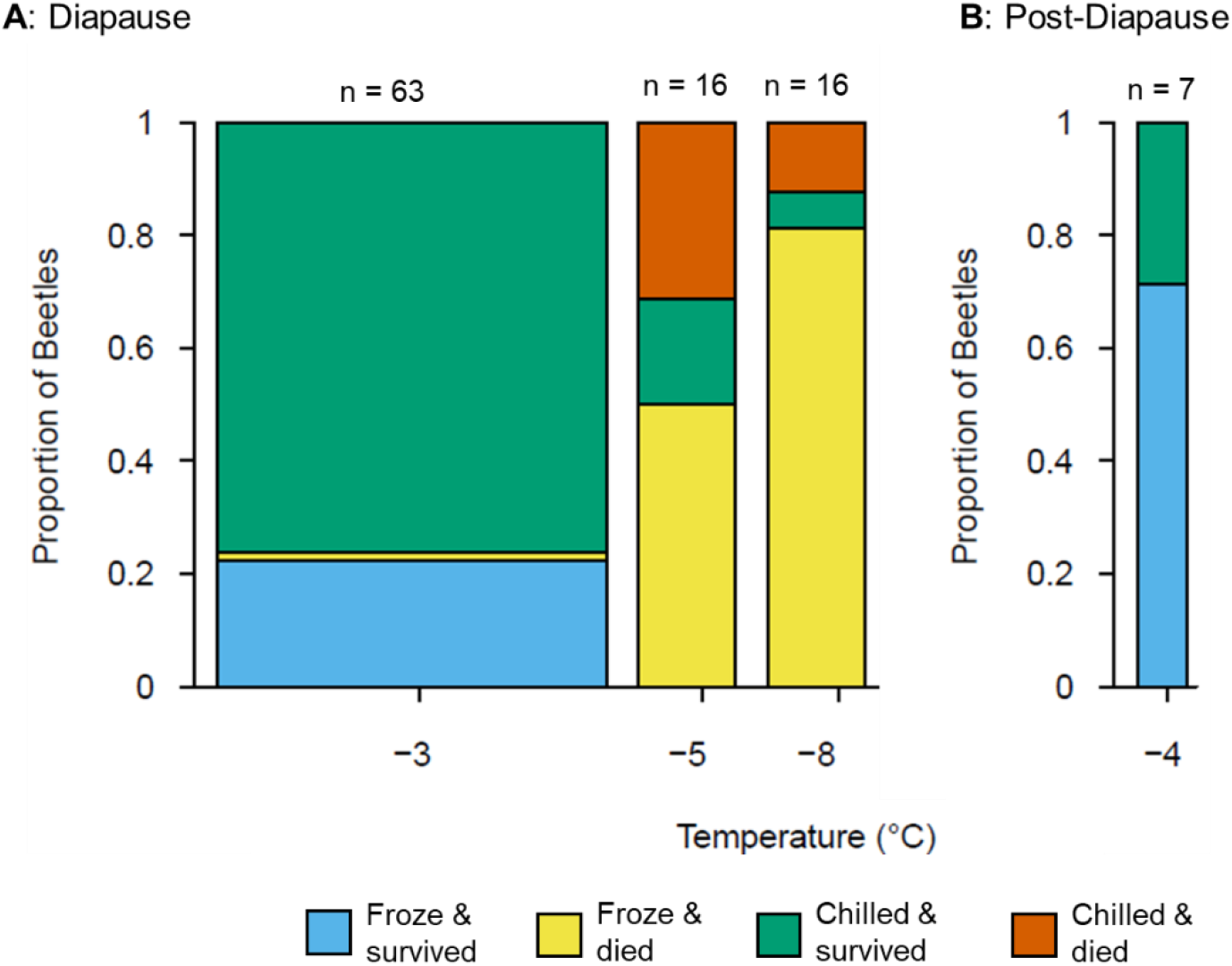
Survival of *Hippodamia convergens* following chilling and freezing at temperatures above the supercooling point. Column width scales with sample size (n). **(A)** Freezing was induced in diapause male and female beetles by external contact with ice at −0.5°C before gradual cooling to the target temperature, followed by 2 days recovery at 4°C. **(B)** Post-diapause male beetles were permitted to feed prior to exposure to −4°C for 4 hours (no cooling/warming ramp), followed by 2 days recovery at 4°C.

We note there was some apparent resistance to inoculative freezing at higher subzero temperatures in diapause beetles. For example, for the −3°C treatment, 35 beetles were in contact with ice, but only 15 of them froze (Table 1; Figure 2A). Higher proportions of beetles froze at the lower temperatures (Table 1; Figure 2A; G test: *G*_6_ = 88.632, *P* < 0.001). A high proportion (c. 70%) of post-diapause males froze at −4°C without exposure to external ice or environmental moisture that could freeze (Figure 2B), but we note these beetles had recently been fed and likely had material in their guts that could induce freezing.

### Post-diapause lady beetles can exhibit normal mating behaviours following chilling or freezing

Some post-diapause male beetles that were chilled or frozen at mild low temperatures (0°C or −4°C) for 4 hours engaged in mating within the 3 days following cold exposure, similar to control beetles (Table 2). Over half (18/30) of the beetles observed in our experiments exhibited mating behaviour on at least one of these 3 days (Table 2). Almost all of these beetles (17) first copulated on day 1, while one beetle (in the 0°C group) first copulated on day 2. Within a 4 hour observation period, there was no effect of mating date (day 1, 2, or 3) or exposure temperature on the proportion of males that mated (Table 3; G test: *G*_4_ = 0.083, *P* = 0.999). Therefore, the trend towards a lower proportion of males engaging in copulation after exposure to lower temperatures (Table 3) was not significant.

**Table 3.** Proportion of post-diapause male *Hippodamia convergens* that mated after exposure to 0°C or −4°C for 4 hours in comparison to those that were kept at 4°C (control). Mating was observed at room temperature for 8 hours on day 1, and 4 hours on days 2 and 3 post-cold exposure. Each male was paired with a female that had been maintained at 4°C.

| Pre-mating exposure | Day 1 | Day 2 | Day 3 |
| --- | --- | --- | --- |
| 4°C (control)<sup>a</sup> | 50% (4/8) within 4 hours<br>75% (6/8) within 8 hours | 63% (5/8) | 57% (4/7) |
| 0°C | 20% (3/15) within 4 hours<br>53% (8/15) within 8 hours | 40% (6/15) | 27% (4/15) |
| -4°C<sup>b</sup> | 29% (2/7) within 4 hours<br>43% (3/7) within 8 hours | 29% (2/7) | 14% (1/7) |

<sup>a</sup> One beetle died between the day 2 and day 3 mating trials. This beetle did not mate on any day.
<sup>b</sup> 5/7 of the -4°C beetles froze. One frozen beetle mated on all three days. One frozen beetle mated on day 1 only (within 8 hours).

The time for post-diapause males to initiate mating and the duration of copulation were unaffected by acute low temperature exposures. Over an 8-hour observation period 1 day post-cold exposure, latency ranged from 30 minutes to 7 hours and 35 minutes, with no difference in mean latency among control beetles and beetles exposed to 0°C or −4°C (Figure 3A; ANOVA: *F_2,14_* = 0.565, *P* = 0.581). Of the 17 copulation events during this observation, three were shorter than 30 minutes (1 control, 2 from 0°C group) and were thus considered failed mating attempts. The 14 successful copulations ranged in duration from 30 minutes to 7 hours and 50 minutes, and there was no difference in mean copulation duration among control beetles and beetles exposed to 0°C or −4°C (Figure 3B; ANOVA: *F_2,14_* = 0.087, *P* = 0.917).

**Figure 3.**
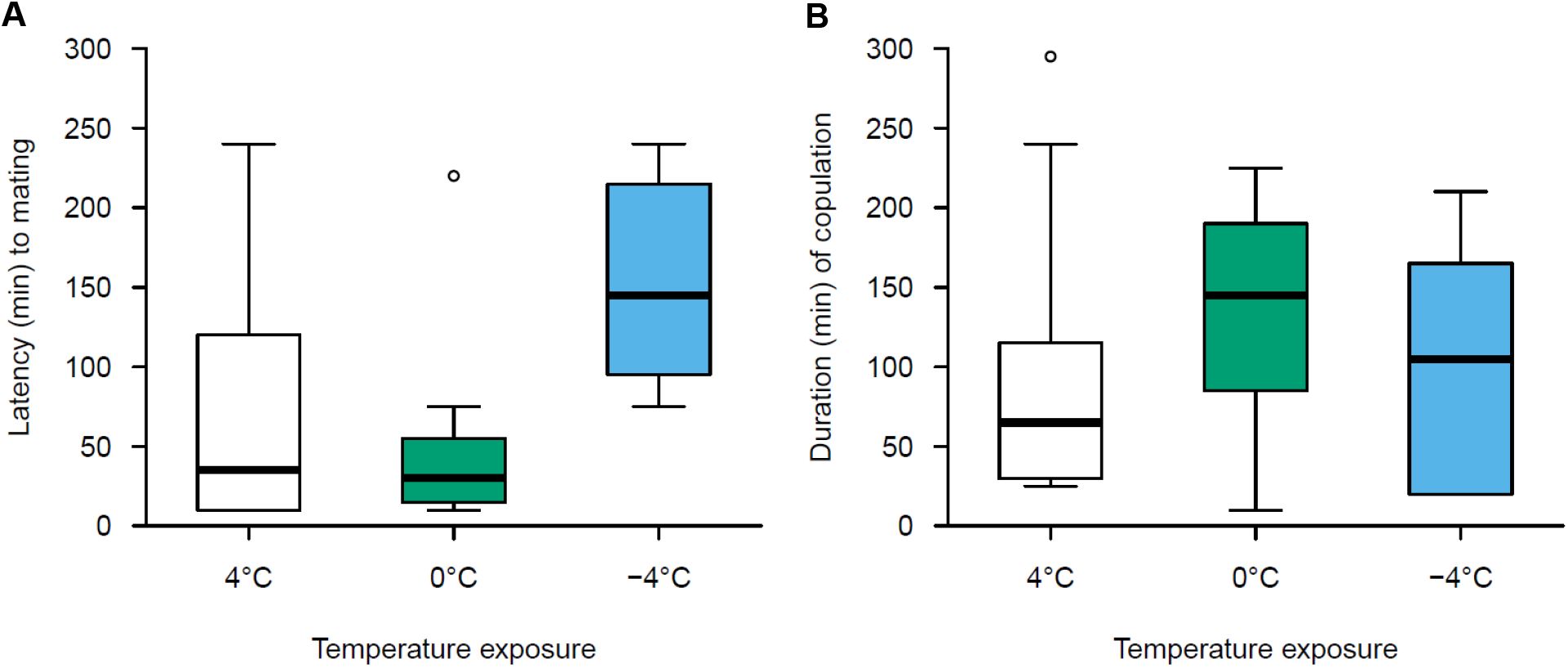
**(A)** Time to initiate mating (latency), and **(B)** duration of copulation of post-diapause male *Hippodamia convergens* following exposure to low temperatures. Males were exposed to 0°C or −4°C for 4 hours or kept at 4°C (control) and then one day post-cold exposure were paired with a female that had been maintained at 4°C. Mating was observed at room temperature for 8 hours. Sample sizes (number of mated males) are in Table 2; within the −4°C group two beetles were frozen and one remained chilled. The thick central line is the median, with the lower and upper bounds of the box representing the first and third quartile, respectively. Whiskers extend to 1.5 times the interquartile range or the most extreme data point above/below the box, whichever is shortest.

When comparing mating behaviours during 4-hour observation periods 1, 2, or 3 days post-cold exposure, there was no effect of observation date on latency (Two-way ANOVA; Treatment: *F_2,28_* = 4.262, *P* = 0.027; Observation Day: *F_2,26_* = 2.491, *P* = 0.106; Treatment × Observation Day: *F_4,22_* = 0.658, *P* = 0.628), so the data are not visualized. There was an effect of observation date on copulation duration, which decreased on later observation dates in all treatment groups (Two-way ANOVA; Treatment: *F_2,28_* = 1.810, *P* = 0.187; Observation Day: *F_2,26_* = 6.956, *P* = 0.005; Treatment × Observation Day: *F_4,22_* = 1.053, *P* = 0.403).

### Diapause lady beetles accumulate a variety of cryoprotectants

We detected several putative cryoprotectant molecules in fat body tissue from diapause *H. convergens*. These cryoprotectants included glycerol, *myo*-inositol, proline, and trehalose, with glycerol and proline being the most abundant (Figure 4). We also tested for mannitol and sorbitol, but they were not present at detectable levels in our assays (data not shown). Exposure to −3°C for 20 hours did not alter cryoprotectant abundance (Figure 4), even if beetles were frozen. There was no effect of treatment on concentrations of glycerol (ANOVA: *F*_2,41_ = 0.418, *P* = 0.661), *myo*-inositol (ANOVA: *F*_2,43_ = 0.508, *P* = 0.605), proline (ANOVA: *F*_2,44_ = 0.169, *P* = 0.846), or trehalose (ANOVA: *F*_2,41_ = 0.217, *P* = 0.806), although there was a trend towards higher glycerol in beetles that were frozen rather than chilled at −3°C (Figure 4).

**Figure 4.**
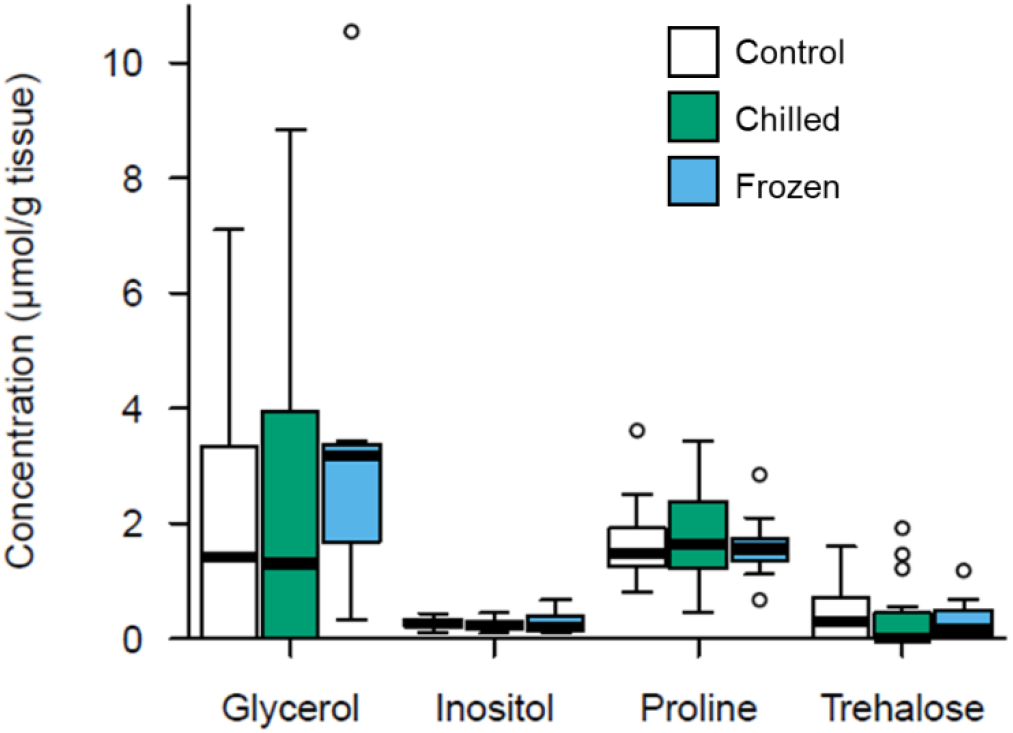
Cryoprotectant concentrations in fat body sampled from *Hippodamia convergens* following no treatment (control; maintained at 4°C) or exposure to −3°C for 20 h in a frozen or unfrozen state followed by 2 days recovery at 4°C. Freezing was induced by external contact with ice at −0.5°C before gradual cooling to the target temperature. The thick central line is the median, with the lower and upper bounds of the box representing the first and third quartile, respectively. Whiskers extend to 1.5 times the interquartile range or the most extreme data point above/below the box, whichever is shortest.

## Discussion

Our work documents the first example of a lady beetle that survives freezing. When cold tolerance strategy was determined via traditional methods (Sinclair et al., 2015) and in the absence of ice nucleators, *H. convergens* survived low temperatures by freeze avoidance. However, when freezing was either inoculated by external contact with ice (Rozsypal and Koštál, 2018) or by nucleated by gut contents at relatively high subzero temperatures (−3 or −4°C), the beetles were freeze tolerant. Contrary to our prediction, males were able to engage in normal mating behaviour after a short freezing event (4 hours at −4°C), suggesting no negative effects of freezing on neuromuscular and endocrine system function. In addition, we detected several putative cryoprotectants (glycerol, proline, *myo*-inositol, and trehalose) in *H. convergens* fat body tissue that may support their cold tolerance, although none of them had a statistically significant increase in concentration following mild chilling or freezing at −3°C for 20 hours - contrary to our prediction. This initial characterization of *H. convergens* freeze tolerance lays the groundwork for future investigations of the mechanisms underlying inoculative freeze tolerance in this beneficial insect.

Hippodamia convergens *can tolerate freezing at high subzero temperatures*

The ability of *H. convergens* to survive inoculative freezing is similar to the firebug *Pyrrhocoris apterus*, which is typically classified as freeze-avoidant (or freeze-intolerant) but is freeze-tolerant when inoculatively frozen by contact with external ice at high subzero temperatures (e.g., −2.5°C) for durations similar to our experiment (Rozsypal and Koštál, 2018). Thus, although prior work describes *H. convergens* as freeze-intolerant (Bennett and Lee, 1989; Lee, 1980), we show that they can survive freezing under mild conditions such as exposure to −3°C or −4°C, and that gut contents can potentially stimulate survivable freezing in post-diapause beetles. In addition, higher proportions of beetles were inoculatively frozen at lower temperatures (43% at −3°C vs. 87% at −8°C; Table 1), suggesting it becomes more challenging to resist inoculative freezing by contact with external ice as temperatures decrease. Freeze tolerance may be adaptative for *H. convergens* in their natural environment (Toxopeus and Sinclair, 2018), where freezing could be stimulated in winter by contact with frost or snow, or in spring (when subzero cold snaps are still possible) by gut contents associated with feeding.

Our *H. convergens* supercooled to approximately −15°C and survived brief exposures to temperatures close to −15°C as long as they did not freeze, which is consistent with previous work on these beetles (Bennett and Lee, 1989). Freeze avoidance via supercooling point depression is a common cold tolerance strategy among lady beetles, including *H. variegata* (Hamedi and Moharramipour, 2013; Khabir et al., 2023), *Hippodamia quinquesignata* (Harper and Lilly, 1982), and *H. axyridis* (Berkvens et al., 2010; Koch et al., 2004). In addition, partial freeze tolerance has been described for *H. axyridis* – that is, some beetles survive if cooled to the supercooling point but are removed from the cold after a very short duration of ice formation (Koch et al., 2004). This partial freeze tolerance at the supercooling point is similar to what we observed in a small sample size of *H. convergens* during our cold tolerance strategy experiment, and we thus suggest it would be useful for future studies to test for inoculative freeze tolerance in other lady beetle species with partial freeze tolerance.

Although *H. convergens* could survive freezing to high subzero temperatures (e.g., −3°C) and supercooling to lower subzero temperatures (e.g., −15°C), their overall cold tolerance was somewhat weak. Almost all beetles survived 20 hours at −3°C – regardless of whether they froze or remained unfrozen – and it would be interesting to test how long they can survive these subzero temperatures (i.e., determine lethal time; Sinclair et al., 2015) in future. However, almost no beetles (frozen or unfrozen) survived a 20-hour exposure to −8°C. This suggests that *H. convergens* may die at temperatures well above their supercooling point (−15°C) if exposed to moderate subzero temperatures for longer durations. It is possible that additional acclimation (e.g., rapid cold hardening, fall-like conditions) could enhance cold tolerance in *H. convergens*, similar to other insects (Berkvens et al., 2010; Harper and Lilly, 1982; Teets and Denlinger, 2013; Toxopeus et al., 2019b), allowing them to survive lower subzero temperatures in their natural overwintering environment.

### Mating behaviour is normal after chilling or freezing at high subzero temperatures

There was a trend towards a higher proportion of mating among males exposed to milder temperatures (0°C or 4°C) than those that were exposed to −4°C, but a larger sample size or longer cold exposures (e.g., Sakaki et al., 2019) are likely needed to determine if freezing is indeed detrimental for mating. While our sample sizes for the mating behaviour experiments were small, it is notable that 2 of the 5 males that froze engaged in successful (longer than 30 minutes) copulations one day post-thaw. The ability to reproduce post-freeze has been documented in other freeze-tolerant insects: frozen and thawed pre-pupae of the goldenrod gallfly *Eurosta solidaginis* can molt into adults and lay eggs (Irwin and Lee, 2000; Marshall and Sinclair, 2018), and a small portion of freeze-tolerant *D. melanogaster* larvae can survive post-thaw to the adult stage and produce viable offspring (Koštál et al., 2012). However, studies that freeze adult insects and subsequently investigate reproduction are rare, despite knowledge of multiple freeze-tolerant adults (e.g., *Hemideina maori*, *Sigaus australis*, *Celatoblatta quinquemaculata*; Morgan-Richards et al., 2023). Our study provides a first step towards describing the effect of freezing on mating in adult *H. convergens*, with further work needed to investigate how freezing affects females and other reproductive metrics for both sexes, such as male ability to mate with resisting females (Perry et al., 2009), time to oviposition (Sakaki et al., 2019; Yan et al., 2025) and fecundity (Chang et al., 1996).

Contrary to our predictions, latency to first mating and copulation duration were similar among males exposed to all temperature treatments (4, 0, or −4°C), although a larger sample size is needed to determine whether freezing had a detrimental effect. Copulation durations one day post-thaw were similar to what has been observed in *H. variegata* under non-stressful thermal conditions (Pervez and Singh, 2013), suggesting *H. convergens* had successfully recovered from any potential injury before mating trials began. To our knowledge, latency to first mating hasn’t been described for *H. convergens* (e.g., Bayoumy and Michaud, 2014; Michaud and Qureshi, 2006; Stowe et al., 2021); however, short latency times are associated with higher fitness in other lady beetles (Bista and Omkar, 2015). If freezing does cause damage to *H. convergens*, then future studies of post-mating behaviour in *H. convergens* may show that freezing increases latency to first mating.

### Cryoprotectant accumulation may support overwintering survival

The metabolites we detected in the fat body of *H. convergens* included common cryoprotectants such as glycerol, proline, *myo*-inositol, and trehalose, but we did not detect sorbitol or mannitol. Our concentrations (less than 10 µmol/g wet mass) were generally low or moderate, consistent with earlier work on *H. convergens* showing that none of glycerol, sorbitol, trehalose, or mannitol were present in tissue homogenates at concentrations greater than 1 mM (Bennett and Lee, 1989). Our results are also similar to studies on overwintering *H. variegata* collected in Iran, which had concentrations of *myo*-inositol, glucose, glycerol, and trehalose on a similar order of magnitude (e.g., 5 – 10 µmol/g wet mass) to our study (Hamedi et al., 2013; Khabir et al., 2023). This suggests the cryoprotectants we (and others) have detected are at too low of a concentration to contribute substantially to supercooling point depression, and it is worth investigating other mechanisms that could do so, such as antifreeze proteins (Han and Bauce, 1995; Lee, 2010; Tyshenko et al., 1997). However, moderate cryoprotectant concentrations are associated with freeze tolerance: freeze-tolerant *G. veletis* accumulate *myo*-inositol and proline (10 µmol/g wet mass) and trehalose (25 µmol/g wet mass) in their fat body (Toxopeus et al., 2019a). Therefore, the proline and glycerol we detected in *H. convergens* were accumulated to a concentration that may be more important for surviving freezing rather than supporting freeze avoidance.

The low concentrations of putative cryoprotectants we measured in *H. convergens* may be caused by the beetles being kept in captivity and lacking their natural seasonal cues. For example, *H. quinquesignata* in laboratory colonies accumulated no detectable glycerol, but field-collected beetles from Alberta (Canada) did, with higher concentrations in January compared to September or April (Harper and Lilly, 1982). Field-collected *H. axyridis* in Japan accumulated *myo*-inositol to approximately 12,000 µg/g wet mass (Watanabe, 2013), three orders of magnitude higher than our values in *H. convergens*. While our laboratory manipulation of a 20-hour exposure to −3°C did not cause an increase in cryoprotectant concentrations, it would be worth investigating whether exposure to more natural fall-like conditions or lower temperature exposures could simulate substantial accumulation of cryoprotectants such as glycerol.

## Conclusion

Overall, we show that *H. convergens* survives freezing at mild subzero temperatures, which may be a more common trait in freeze-avoidant species than previously thought. The minimal effects of chilling and freezing on male mating behaviours were unexpected, but follow-up studies with larger samples sizes are needed to fully characterize the effects of low temperatures on reproduction. It would be interesting to test the accumulation of the cryoprotectants we detected (glycerol, *myo*-inositol, proline, trehalose) under more natural winter conditions to assess whether they contribute to cold tolerance in *H. convergens*.

## Author Contributions

All authors contributed to experimental design and conceptualization, as well as revising the manuscript. In addition: HEE: mating experiments data collection, analysis; TAK, LEE, MRMW: cold tolerance strategy data collection, analysis; ELB, LJK, JKM, ABS: inoculative freeze tolerance and supercooling point data collection, analysis, MFAR, BRA, AMAR, RPC, SDF, GMG, KEDH, TAK, KEM, JMN, LAP, MKP, ZMS: cryoprotectant data collection, analysis: JCP, JT: supervision, funding, manuscript draft.

## Data Availability Statement

All data and analysis code are available at https://github.com/jtoxopeus/ladybeetles

## Ethics Statement

Ethics Approval was not necessary for this study.

## Acknowledgements

Thank you to L.M. Rogers for assisting with reagent preparation, and the St. Francis Xavier University students in Biology 395 (winter 2025) for piloting the cryoprotectant assays.

